# Fast, scalable prediction of deleterious noncoding variants from functional and population genomic data

**DOI:** 10.1101/069682

**Authors:** Yi-Fei Huang, Brad Gulko, Adam Siepel

**Affiliations:** Simons Center for Quantitative Biology, Cold Spring Harbor Laboratory, Cold Spring Harbor, NY 11724, USA; Graduate Field of Computer Science, Cornell University, Ithaca, NY 14853, USA

## Abstract

Across many species, a large fraction of genetic variants that influence phenotypes of interest is located outside of protein-coding genes, yet existing methods for identifying such variants have poor predictive power. Here, we introduce a new computational method, called LINSIGHT, that substantially improves the prediction of noncoding nucleotide sites at which mutations are likely to have deleterious fitness consequences, and which therefore are likely to be phenotypically important. LINSIGHT combines a simple neural network for functional genomic data with a probabilistic model of molecular evolution. The method is fast and highly scalable, enabling it to exploit the “Big Data” available in modern genomics. We show that LINSIGHT outperforms the best available methods in identifying human noncoding variants associated with inherited diseases. In addition, we apply LINSIGHT to an atlas of human enhancers and show that the fitness consequences at enhancers depend on cell-type, tissue specificity, and constraints at associated promoters.

## Introduction

In the human genome, a large majority of nucleotides that are associated with diseases or other phenotypes, or that show signatures of natural selection, falls outside of protein-coding genes^1–3^. Many of these nucleotides appear to fall in *cis*-regulatory elements, including promoters, enhancers, and insulators. Similar observations hold across most animals and plants, as well as some other eukaryotes^4–7^. Nevertheless, the capability to identify and characterize functionally important noncoding sequences is decades behind that for protein-coding sequences. Investigators still lack a deep understanding of many fundamental properties of these sequences, including the manner in which they interact with chromatin and transcription factors and the biophysical dynamics of protein / nucleic acid complexes important in transcriptional and posttranscriptional regulation. This limited ability to make sense of noncoding DNA is a major barrier for progress in establishing the genetic basis for many diseases and other phenotypes, with major implications for biomedicine, agriculture, synthetic biology, and other fields.

For more than a decade, the genomics community has pushed for a deeper understanding of the noncoding genomes of many animal and plant species through systematic interrogation with high-throughput biochemical assays for features such as transcriptional activity, chromatin accessibility, and specific histone modifications and transcription factor binding events^8–12^. These efforts have produced a wealth of data for dozens of cell types across a range of organisms, and have helped both to identify many predicted regulatory elements, and to clarify many general aspects of gene regulation. Nevertheless, a substantial gap remains between the outcomes of these high-throughput experiments and a detailed understanding of noncoding function, for several reasons. First, these assays generally measure genomic and epigenomic features roughly correlated with, but not directly indicative of, regulatory function. Second, they generally have relatively low resolution along the genome, identifying regions hundreds of nucleotides long, rather than pinpointing single nucleotides. Third, these measures are highly condition-specific, and data has only been generated for a small subset of cell types and conditions.

As a consequence, there is a pressing need for computational methods that more precisely predict regulatory function by jointly considering the results of numerous such assays together with complementary data, such as annotations of protein-coding genes and measures of evolutionary conservation across species. The development of statistical and machine-learning methods that attempt to address this integrative prediction challenge has emerged as an active, fast-moving area of research. Recently published methods in this area can be roughly divided into three categories: (1) machine-learning classifiers that attempt to separate known disease variants from putatively benign variants using a variety of genomic features (e.g., GWAVA^13^ and FATHMM-MKL^14^); (2) sequence- and motif-based predictors for the impact of noncoding variants on cell-type-specific molecular phenotypes, such as chromatin accessibility or histone modifications (e.g., DeepBind^15^, DeepSEA^16^ and Basset^17^); and (3) evolutionary methods that consider data on genetic variation together with functional genomic data and aim to predict the effects of noncoding variants on fitness (e.g., CADD^18^, DANN^19^, FunSeq2^20^, and fitCons^3^). A limitation of the first two classes of methods is that they depend strongly on the available training data, which may be limited and may not be representative of the broader class of regulatory sequences of interest. By contrast, the evolutionary methods obtain their signal not primarily from previously assigned class labels, but instead from signatures of natural selection across the genome over many generations, and they are therefore much less data limited. This approach is likely to be particularly powerful for regulatory sequences that tend to be under strong purifying selection, such as Mendelian disease variants. Evolution-based methods also naturally integrate over cell-types, an important strength when the relevant tissue- or cell-types for a condition of interest are unknown (as with many human diseases).

Among the available evolution-based methods, fitCons is unique in explicitly characterizing the influence of natural selection at each genomic site of interest using a full probabilistic evolutionary model and patterns of genetic variation within and between species. FitCons makes a distinction between functional genomic and comparative genomic data, first defining several hundred clusters of genomic positions with distinct functional genomic “signatures,” and then estimating the fraction of nucleotides under natural selection within each cluster from polymorphism and divergence data. These estimates are obtained using the INSIGHT evolutionary model^21, 22^, and are interpreted as the probabilities that mutations in each cluster of genomic sites will have fitness consequences (fitCons scores). In this manner, fitCons aggregates information about natural selection from a large number of sites with similar functional profiles based on evolutionary first principles. FitCons provides a useful, easily interpretable readout along the genome, complementary to conventional evolutionary conservation scores, and it performs well in predicting cell-type-specific functional elements. A major limitation of the method, however, is that it scales poorly with the available functional genomic data. In particular, the number of clusters considered by the method increases exponentially with the number of functional genomic annotations. This exponential dependency limits the method's ability to take advantage of the growing body of available functional genomic data and therefore limits prediction power. A related problem is that the restriction to small numbers of genomic features leads to a relatively coarse-grained, blocky pattern of scores along the genome, which does not allow for fine distinctions among nearby nucleotide sites.

In this paper, we describe a new method, *<u>L</u>inear* <u>INSIGHT</u> (LINSIGHT; pronounced *lin-site*), that is based on the existing INSIGHT/fitCons framework but has vastly improved speed, scalability, genomic resolution, and prediction power. The main idea behind LINSIGHT is to bypass the clustering step of fitCons and instead couple the probabilistic INSIGHT model directly to a generalized linear model for genomic features. This results in a more streamlined model that scales linearly, rather than exponentially, with the available data, and can make direct use of the input data, with no need for discretization. This generalized linear model can be regarded as a simple neural network, and it readily extends to more complex, multi-layered networks, allowing for nonlinearities and interdendencies among genomic features. By integrating a large number of genomic features, LINSIGHT provides a systematic, high resolution description of the fitness consequences of noncoding mutations in the human genome. We demonstrate that LINSIGHT outperforms state-of-the-art prediction methods in the task of prioritizing noncoding disease variants from the Human Gene Mutation database (HGMD)^23^ and the NCBI ClinVar database^24^. Furthermore, we use LINSIGHT to show that the evolutionary constraints on human enhancers depend on their associated tissue types, degree of tissue specificity, and associated promoters, which has important implications for understanding the evolution of *cis*-regulatory elements and for improving variant prioritization methods. Our LINSIGHT scores are available as a track on the Cold Spring Harbor Laboratory mirror of the UCSC Genome Browser (hg19 assembly).

## Results

### LINSIGHT combines INSIGHT with a scalable linear model

The original INSIGHT and fitCons methods^3,21, 22^ infer the selective pressure on noncoding sites, and hence the likely fitness consequences of noncoding mutations, by contrasting patterns of genetic variation at each focal site with the patterns at nearby genomic regions that are likely to be free from the influence of selection (“neutrally evolving sites”). To address the problem that genetic variation within species and between closely related species (such as the human and chimpanzee) are sparse across the genome, fitCons pools information across the thousands of genomic sites assigned to each discrete cluster.

The key idea behind LINSIGHT is instead to accomplish this pooling of information across sites indirectly, using a generalized linear model (Figure 1; see Supplementary Text and Supplementary Table 1 for complete details). In particular, the parameters of the INSIGHT model that describe natural selection (*ρ* and *γ*) are determined as linear-sigmoid functions of the genomic features local to each site. Thus, the fitness consequences of mutations at each site are assumed to depend on genomic features at that site, such as its RNA expression level (RNA-seq read depth), chromatin accessibility (DNase-I hypersensitive sites), histone modifications or bound transcription factors (ChIP-seq peaks), as well as features based on annotations (e.g., distance to nearest transcription start site, match to known TFBS motif) and comparative genomics (e.g., phyloP or phastCons scores). This approach has several major advantages: it requires no clustering and no discretization *a priori*, and it scales linearly with the available genomic features, allowing hundreds or even thousands of features to be considered. In contrast to fitCons, the scalability of the method enables data to be pooled across cell-types, and it allows the scores to reach single-nucleotide resolution along the genome. Nevertheless, LINSIGHT continues to benefit from the advantages of the probabilistic INSIGHT model of molecular evolution.

**Figure 1:**
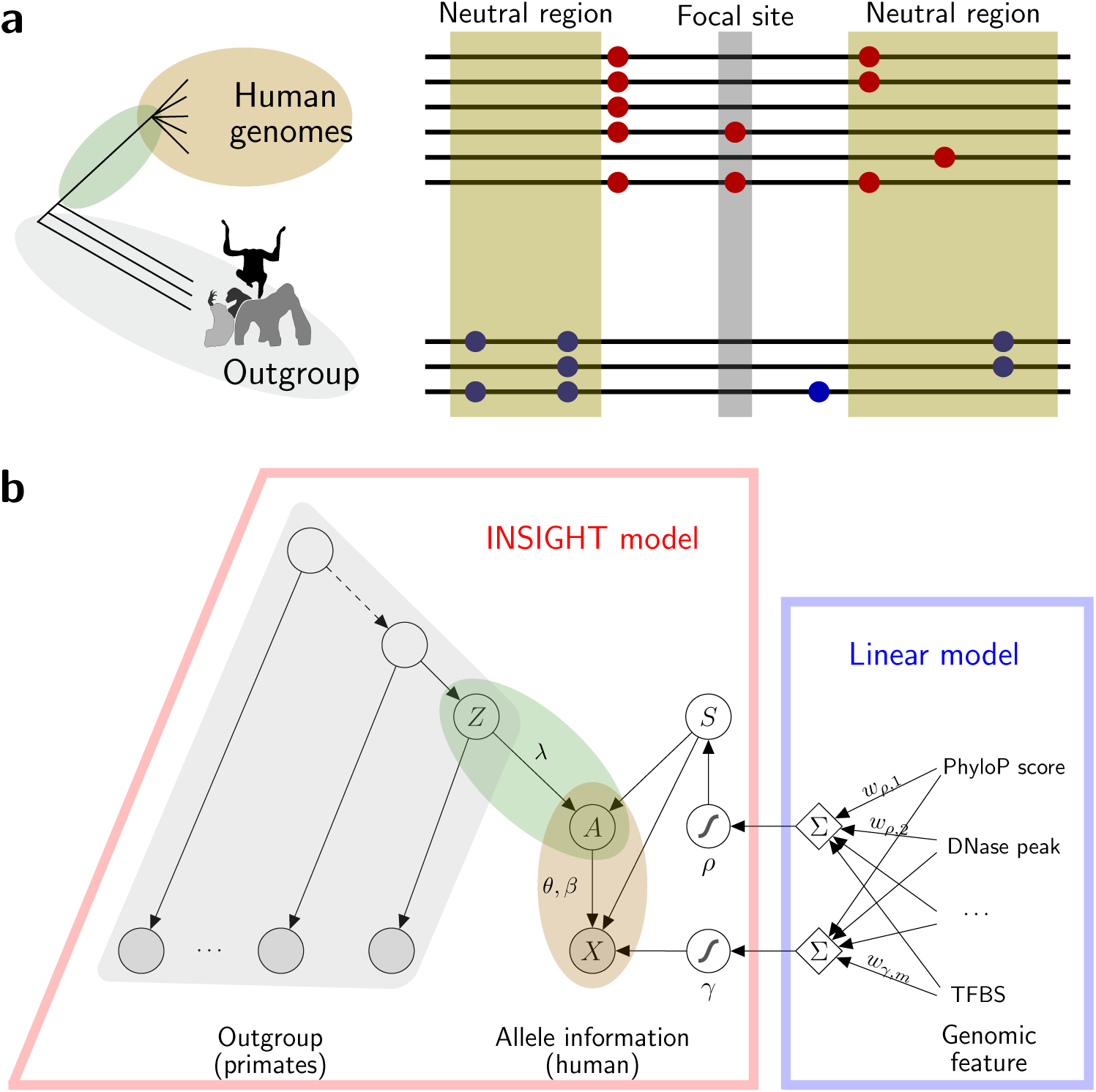
Conceptual overview of LINSIGHT. **(a)** Like our previous fitCons^3^ method, LINSIGHT estimates probabilities that mutations at each genomic site will have fitness consequencs, based on patterns of genetic polymorphism within a species (here, humans) and patterns of divergence from closely related outgroup species (chimpanzee, orangutan, and rhesus macaque). Patterns of genetic variation at the focal site and other sites like it are contrasted with those in neutrally evolving regions nearby. Red circles indicate human single nucleotide polymorphisms and blue circles indicate nucleotide substitutions between species. **(b)** LINSIGHT combines the probabilistic graphical model from INSIGHT ^21,22^ with a generalized linear model. The selection parameters from INSIGHT, *ρ* and *γ*, are defined by linear combinations of local genomic features, followed by sigmoid transformations. *ρ*: probability that focal site is under negative selection (hence, has fitness consequences if mutated); *γ*: relative rate of low frequency polymorphism for focal sites under selection; *S*: indicator of natural selection; *Z*: ancestral allele in the human-chimpanzee most recent common ancestor (MRCA); *A*: ancestral allele in the MRCA of all modern humans; *X*: allelic state in human population, including major and minor alleles and minor allele frequency; *w_ρ,i_*: weight of genomic feature *i* for parameter *ρ*; *w_γ,i_*: weight of genomic feature *i* for parameter *γ*. *λ*: block-wise neutral divergence rate. *θ*: block-wise polymorphism rate. *β* = (*β*_1_, *β*_2_, *β*_3_): proportions of low, intermediate, and high frequency neutral derived alleles genome-wide. Note that a third selection parameter from INSIGHT, *η*, is omitted here, because positive selection has a negligible effect in this setting (see Supplementary Text).

All parameters of the LINSIGHT model can be estimated simultaneously from genome-wide data by maximum likelihood using an online stochastic gradient descent algorithm (Methods). The gradients for the feature weights can be efficiently computed by the back-propagation method widely used in neural network training^25^. Indeed, the model can be considered a type of neural network, albeit one without hidden layers. Its main disadvantage relative to fitCons—the assumption of an additive, linear relationship between features and selection parameters—could be addressed by adding hidden layers to the neural network, although we have found its performance to be excellent without this extension. Notably, the amount of data available for training is large in comparison to the number of free parameters and we have not found regularization to be necessary, but it could easily be added if necessary.

### LINSIGHT scores across the human genome are generally consistent with, but often improve on, previous measures of evolutionary conservation

We applied LINSIGHT to a large public data set consisting of complete genome sequences for multiple human individuals and nonhuman primates, comparative genomic data for mammals and vertebrates, and a wide variety of functional genomic data, and we generated LINSIGHT scores for all positions across human reference genome (Methods). We considered a total of 48 genomic features, falling in three general classes: conservation scores, predicted binding sites, and regional annotations (Table 1 and Supplementary Table 2). We used the human polymorphism data (from the Complete Genomics “69 Genomes” data set) and primate divergence data (from alignments of the human, chimpanzee, orangutan, and rhesus macaque genomes) that were used for fitCons^3^. Note that, while it might appear circular to use evolutionary conservation scores as features since LINSIGHT's objective function is also essentially a measure of conservation, these scores reflect the influence of natural selection over alternative time scales (e.g., ∼80 million years of mammalian evolution), and considering them in this way substantially improves prediction performance (as shown below; see Discussion).

**Table 1:**
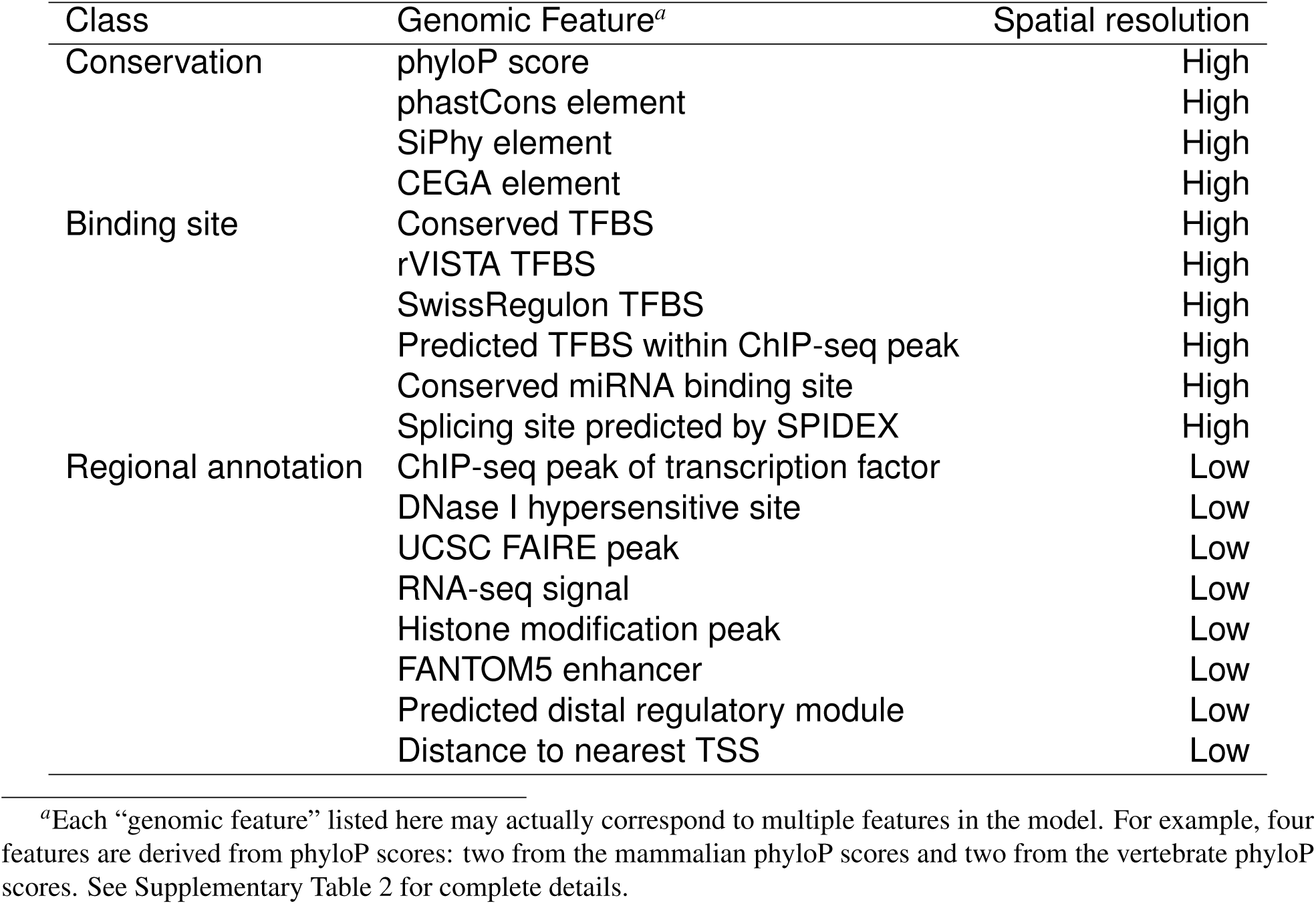
Summary of genomic features used for LINSIGHT scores

The distributions of INSIGHT scores in annotated regions of the noncoding genome are generally consistent with previous observations based on conservation scores^1,4,26^. For example, splice sites are very highly constrained (median LINSIGHT score of 0.956, indicating a 95.6% probability of fitness consequences due to mutations at these nucleotide sites), whereas annotated TFBSs show reduced, but still substantial, constraint (median score of 0.240 for TFBSs shared across species, median score of 0.106 for all TFBSs from the Ensembl Regulatory Build^27^; Figure 2a). Other promoter regions (median score of 0.073) and untranslated regions (UTRs; median scores of 0.128 and 0.076 for 5′ and 3′ UTRs, respectively) are somewhat less constrained, and unannotated intronic and intergenic regions exhibit the least constraint (median scores of 0.044–0.048). As observed previously, 5′ UTRs show somewhat more constraint than 3′ UTRs, although both types of UTRs contain subsets of sites subject to strong selection (LINSIGHT score >0.8)^4,26^. The estimate for the more conserved TFBSs (0.240) is roughly similar to, if slightly lower than, previous estimates directly obtained from experimentally defined TFBSs (∼30-40% of sites under selection^22,28^), de-spite that it was obtained indirectly in this case via the generalized linear model. The genome-wide average of the LINSIGHT scores is about 0.07, suggesting that about 7% of noncoding sites are under evolutionary constraint, consistent with numerous previous studies^3,4, 29–31^.

**Figure 2:**
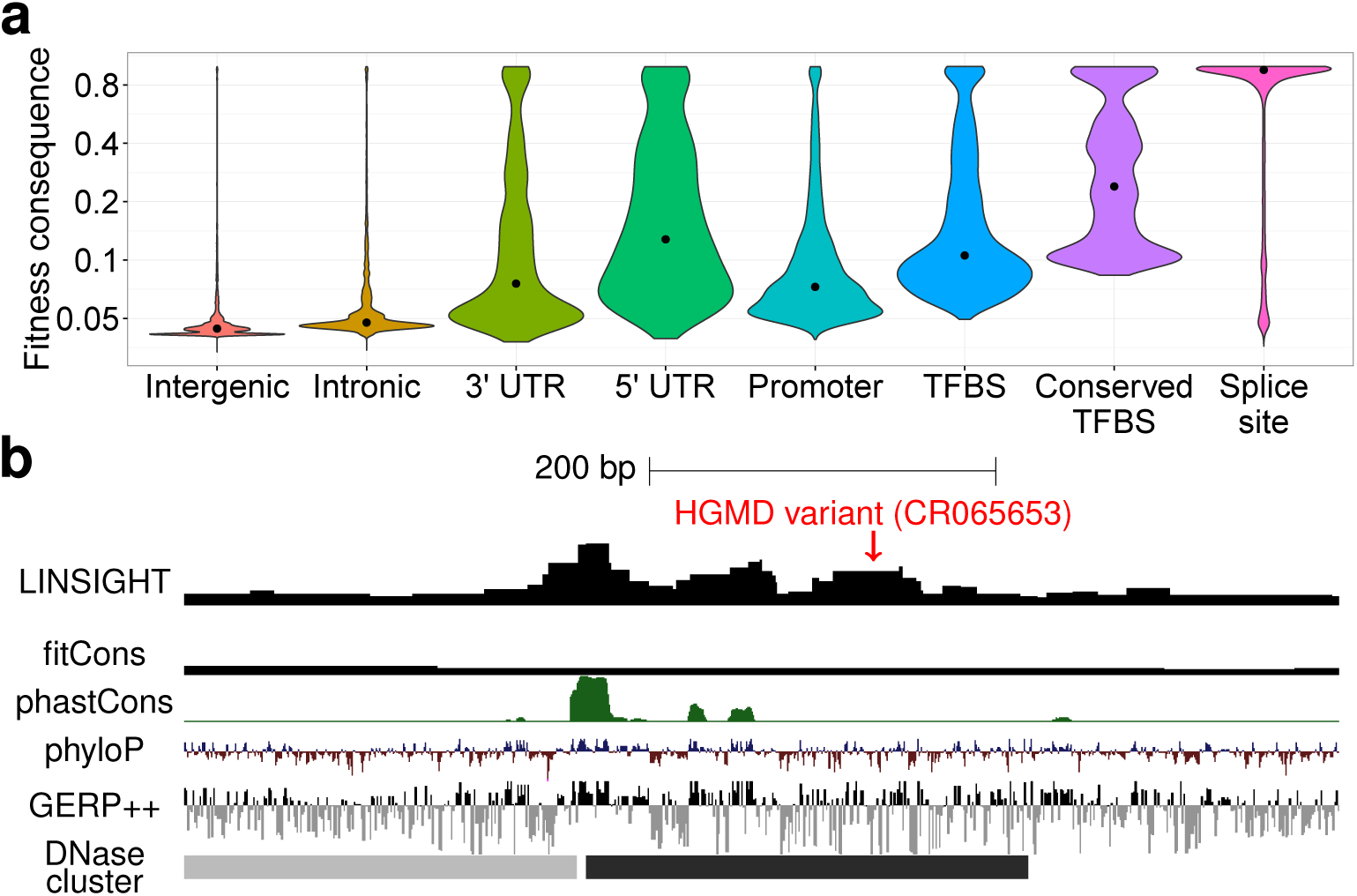
Summary of LINSIGHT scores across the human genome. **(a)** Distributions of LINSIGHT scores for various genomic regions. Intergenic, intronic, UTRs, and 1-kb promoters were defined based on GENCODE annotations; TFBSs were predicted from ChIP-seq peaks (Ensembl Regulatory Build); and conserved TFBSs were obtained from the UCSC Genome Browser. Note the logarithmic scale. **(b)** UCSC Genome Browser display showing LINSIGHT scores alongside those from fitCons, phastCons, phyloP, and GERP++. LINSIGHT integrates functional genomic data together with conservation scores and other features to provide a high-powered, high-resolution measure of potential function. In this example, it is the only method to highlight a variant from HGMD (CR065653) that is associated with up-regulation of the telomerase reverse transcriptase (*TERT*) gene.

Across all noncoding positions in the genome, the LINSIGHT scores are fairly well correlated with scores from recently published methods such as FunSeq2^20^ and Eigen^32^ (Spearman's *ρ* = 0.43–0.48), moderately with those from fitCons^3^ (*ρ* = 0.33), and relatively poorly with those from phyloP^26^, GERP++^33^, and CADD^18^ (*ρ* = 0.04–0.06; Supplementary Figure 1a). However, these sitewise correlations are strongly influenced by large regions of the genome bereft of functional genomic data. Within phastCons-predicted conserved elements^4^, which are strongly enriched for regulatory function, LINSIGHT's correlation increases to *ρ* = 0.62–0.66 for phyloP, GERP++, and Eigen, and to *ρ* = 0.43 for CADD, while decreasing to *ρ* = 0.42 for FunSeq2 and to *ρ* = 0.16 for fitCons (Supplementary Figure 1b). In general, the scores that make use of functional genomic data, including those from LINSIGHT, FunSeq2, Eigen, and CADD, are relatively well correlated, as are those from the pure conservation methods, phyloP and GERP++, but these two groups of scores are less well correlated with one another. The fitCons method shows the least correlation with other methods, primarily because of its low genomic resolution.

On the task of identifying likely regulatory elements in unannotated regions of the genome, the functional genomic methods generally perform better than pure conservation methods, and LINSIGHT is among the best available methods at this task. For example, LINSIGHT has good power to identify transcription factor binding sites from the ORegAnno database^34^, with an AUC = 0.926, outperformed only by DeepSEA (AUC = 0.965) and FunSeq2 (AUC = 0.950) among seven methods tested (Supplementary Figure 2). These three methods perform substantially better than conservation-based methods (e.g., phyloP has AUC = 0.884) as well as methods that use functional genomic data such as GWAVA (AUC = 0.814), CADD (AUC = 0.841), and Eigen (AUC = 0.899). Thus, despite that it relies on an evolutionary objective function, LINSIGHT maintains good performance in the prediction of regulatory elements, competitive with methods optimized for regulatory element prediction and superior to methods that consider evolutionary conservation alone.

Consistent with these general trends, LINSIGHT highlights many of the regions identified by conservation-methods such as phastCons, phyloP, and GERP++, but also identifies some regions that have relatively low conservation scores yet are likely to have important biological functions. An example is HGMD variant CR065653, associated with up-regulation of the telomerase reverse transcriptase (*TERT*) gene, which obtains an elevated LINSIGHT score, but is not identified by phastCons, phyloP, or GERP++ as being under constraint (Figure 2b). This example also demonstrates that the genomic resolution of the LINSIGHT scores is dramatically better than that of fitCons, and approaches the nucleotide resolution of phyloP and GERP++.

### LINSIGHT accurately identifies disease-associated variants in noncoding regions

We tested the ability of LINSIGHT to identify noncoding nucleotide positions that are associated with inherited human diseases, using the HGMD^23^ and ClinVar^24^ databases to define positive examples, and common polymorphisms (MAF > 1%), which are unlikely to be functionally important, to define negative examples. For comparison, we evaluated the CADD^18^, Eigen^32^, DeepSEA^16^, FunSeq2^20^, GWAVA^13^, and phyloP^26^ methods on the same task. For each scoring method, we computed false positive vs. true positive rates for the complete range of score thresholds, displaying the results as receiver operating characteristic (ROC) curves and measuring prediction power by the area-under-the-curve (AUC) statistic. Because the results of these tests can be highly sensitive to the criteria for selecting negative examples, we considered three schemes of increasing stringency (following ref. [13]): all negative examples (unmatched), negative examples matched by distance to the nearest transcription start site (matched TSS), and negative examples matched by specific genomic region (matched region; see Methods for details).

Overall, LINSIGHT outperformed all other methods in all comparisons (Figure 3 and Supplementary Figure 3). Its absolute prediction power varied across matching schemes in a predictable manner, being highest in the unmatched comparison (e.g., AUC = 0.897 for HGMD) and decreasing in the matched TSS (AUC = 0.759) and matched region (AUC = 0.660) comparisons. The same effect also occurred for most other methods, but the methods that make heavier use of regional information, such as FunSeq2, suffered more as the matching stringency increased. In almost all cases, the AUCs were considerably higher for ClinVar than for HGMD, apparently because ClinVar is heavily enriched for variants in splice sites, which are relatively easy to identify (Supplementary Figure 4). An exception to this rule was GWAVA, which performs exceptionally well on HGMD (cross-validation AUCs of 0.71–0.97)^13^ and much more poorly on ClinVar (AUCs of 0.741–0.887), but GWAVA was trained using HGMD^13^ and its performance on that data set appears to reflect overfitting (not shown in the ROC plots for this reason). This dependency on the training set for GWAVA demonstrates one of the pitfalls of pure classification strategies, and highlights a strength of the evolution-based strategy, which are much less dependent on a training set. Nevertheless, phyloP performs quite poorly on the HGMD data set, and CADD is only slightly better, showing that scores based exclusively or primarily on evolution are of limited usefulness in this task. The excellent performance of INSIGHT in the tests appears to derive from its use of both a broad collection of informative features along the genome and an evolution-based objective function.

**Figure 3:**
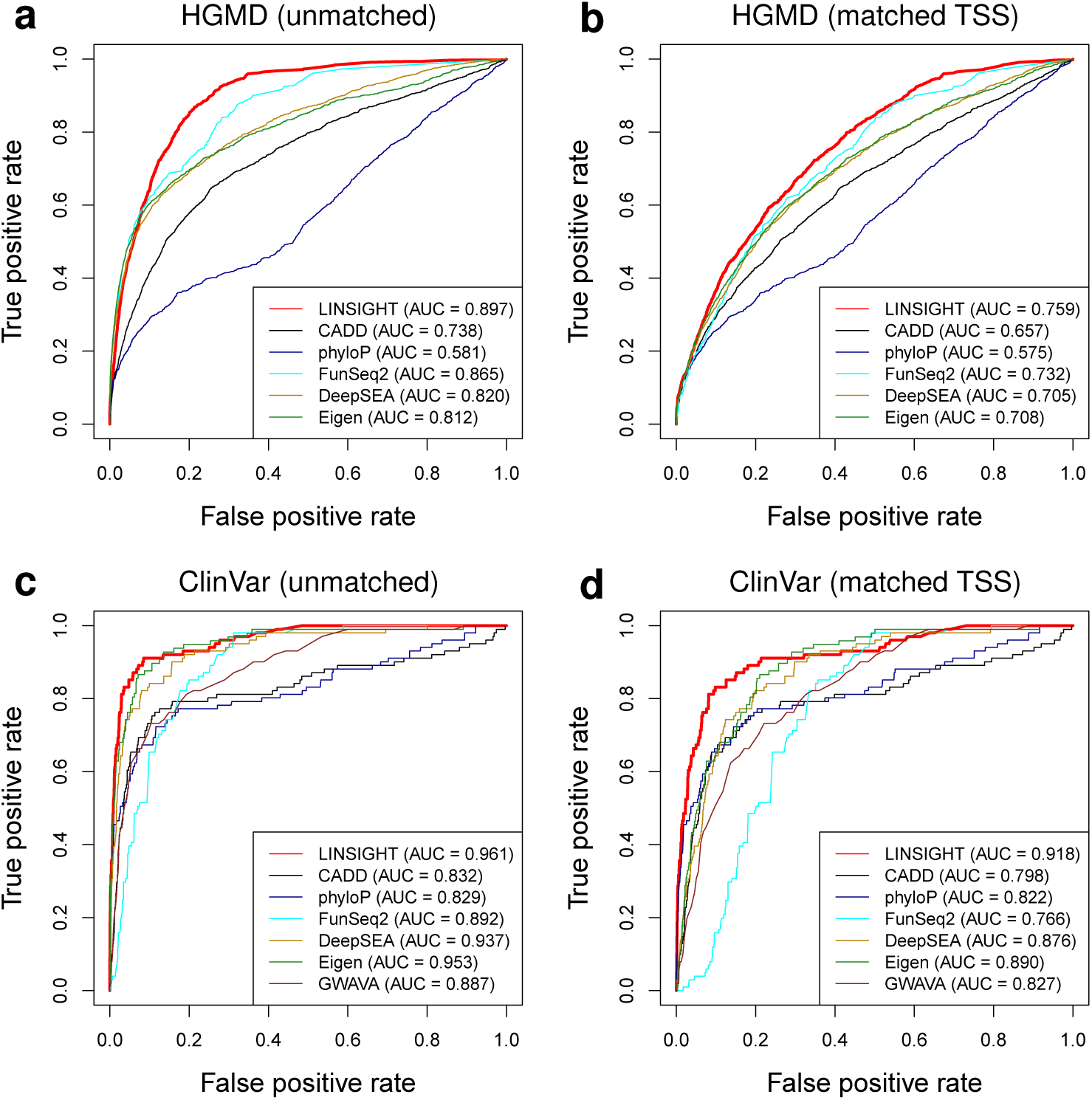
Prediction power of various computational methods for distinguishing disease-associated noncoding variants from variants not likely to have phenotypic effects. Power is described using receiver operating characteristic (ROC) curves and quantified using the Area Under the Curve (AUC) statistic. Results are shown for positive examples from the HGMD^23^ (panels **a** & **b**) and ClinVar^24^ (**c** & **d**) databases. Common SNPs (MAF > 1%) were used as negative examples and were either randomly selected (**a** & **c**) or matched to positive examples by distance to nearest transcription start site (**b** & **d**). LINSIGHT is compared with CADD^18^, phyloP^26^, FunSeq2^20^, DeepSEA^16^, Eigen^32^, and GWAVA^13^. FitCons is not included because it performs poorly on this task due to its low genomic resolution and cell-type specificity. GWAVA is not shown in **(a)** & **(b)** because it was trained on HGMD data.

### The relative contributions of genomic features to LINSIGHT prediction performance are context-dependent

The genomic features used by LINSIGHT can be grouped into three broad classes: conservation scores, predicted binding sites, and regional annotations (Table 1). We examined the relative contributions to prediction power of these feature classes by retraining the model three times, each time removing a different class of features. This procedure was applied at the level of feature classes, rather than individual features, because of the strong correlations among the features within each class. We measured the prediction power of each version of the model using the AUC statistic, as above, but this time we merged the HGMD and ClinVar variants and then divided them into four categories based on their locations relative to genomic annotations: variants in promoters of protein coding genes; variants in 5′ or 3′ UTRs; variants proximal to splice sites; and all other noncoding variants. As a measure of the contribution of each class of features, we used the reduction in AUC resulting from the exclusion of that feature class. Thus, a large reduction in AUC implies that the removed features are highly important in prediction, while a small reduction implies that they are less important.

These feature classes provide complementary information about disease variants, but their relative contributions depend strongly on both the matching scheme for positive and negative examples and on the genomic locations of the variants of interest (Figure 4 and Supplementary Figure 5). For example, regional annotations are powerful predictors when unmatched negative controls are used (Figure 4a), because these features are broadly useful in distinguishing genomic regions enriched for functional variants from the genomic background. However, when matched TSS or matched region controls are considered (Figure 4b and Supplementary Figure 5), regional annotations diminish in importance and, in most genomic regions, conservation scores make a larger contribution to prediction performance. This is because the higher genomic resolution of conservation scores makes them much more useful in distinguishing functional sites from nearby nonfunctional sites. Interestingly, predicted binding sites make a substantial contribution only in promoter regions and in the case of matched controls (Figure 4e and Supplementary Figure 5a), apparently because these binding sites (most of which are TFBSs) are strongly enriched in these regions. Altogether, these results indicate the contributions of genomic features to prediction power for disease-associated noncoding variants depends on both the genomic regions considered and the details of the comparison scheme. This observation may help to explain some of the discrepancies in the literature regarding the apparent relative prediction performance of the available methods.

**Figure 4:**
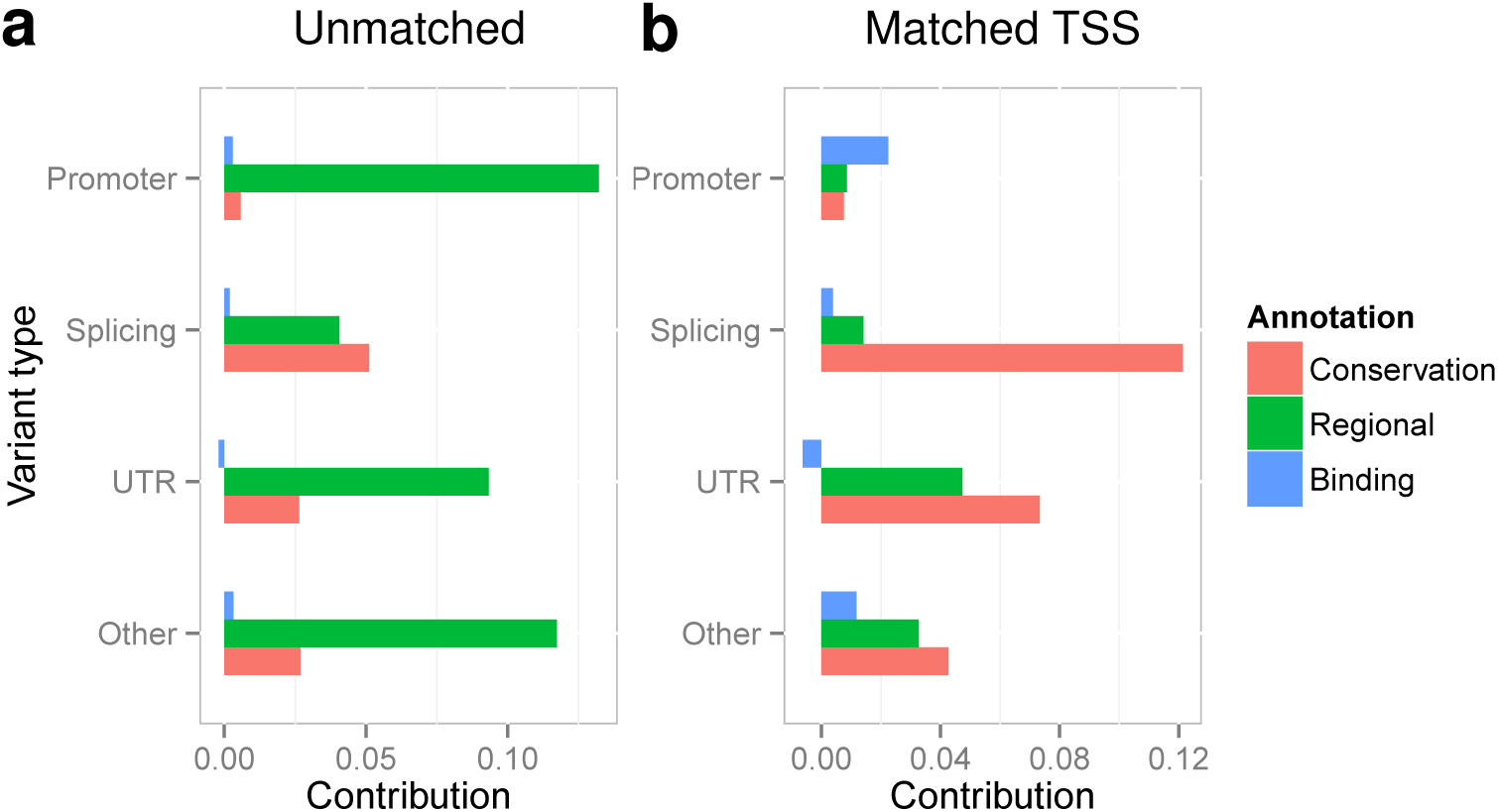
Contributions of various genomic features to the identification of disease-associated variants from HGMD and ClinVar. The contribution of each class of genomic features is measured as the reduction in the area under the curve (AUC) statistic resulting from the removal of those features. Results are shown for four types of variants and two matching schemes for positive and negative examples. Additional results are shown in Supplementary Figure 5.

### The evolutionary constraints on enhancers are context-dependent

In addition to its value in identifying regulatory sequences and predicting disease relevance, LINSIGHT is potentially useful for studying the influence of natural selection on noncoding sequences. Compared with other measures of selection, LINSIGHT has the advantages of considering both functional genomic and population genomic data, of detecting the influence of selection on relatively recent time scales (e.g., since the human/chimpanzee divergence), and of providing a model-based, easily interpretable measure of fitness consequences. With these advantages in mind, we used the method to gauge the degree of evolutionary constraint on enhancers in the human genome, considering in particular the relationships between constraint and the number and type of active cell types and the target promoter of each enhancer. We analyzed nearly 30,000 enhancers (median length 293 bp) from a recently published atlas of active enhancers in dozens of human cell types and tissues, which were identified based on their transcriptional signatures^35^. This approach of annotating enhancers based on enhancer-associated RNAs (eRNAs) has been shown to identify elements having active roles in gene regulation in a cell-type-specific fashion with high genomic resolution^35–37^.

First, we examined the relationship between the LINSIGHT scores and the number of cell types in which each enhancer is active. We found that the LINSIGHT scores were significantly positively correlated with the number of active cell types (*p* < 10^−15^; Figure 5a), indicating that a broader spectrum of activity across cell types is associated with stronger purifying selection. This finding parallels similar findings for protein-coding genes^38–40^ and TFBSs^22^ and likely reflects a general correlation between pleiotropy and constraint (see Discussion). Second, we examined the relationship between the LINSIGHT score for each enhancer and the tissue type in which that enhancer is active, focusing on tissue-specific enhancers (active in a single tissue type). We found that tissue-specific enhancers that were associated with sensory perception (olfactory region and parotid gland), the immune system (lymph node), digestion (stomach), and male reproduction (penis and testis) had the lowest LINSIGHT scores, whereas tissue-specific enhancers associated with tissues such as smooth muscle, the skin, and the urinary tract and bladder had the highest LINSIGHT scores (Supplementary Figure 6). These findings are also broadly consistent with findings for protein-coding genes, which have indicated that sensory, immune, dietary, and male reproductive genes are enriched with positively selected (fast evolving) genes^40,41^. Interestingly, enhancers active in tissues associated with female reproduction (e.g., uterus, female gonad, and vagina) appeared to be under substantially more constraint than those active in tissues associated with male reproduction, perhaps owing to increased positive selection on male reproductive functions. Finally, we compared the LINSIGHT scores at enhancer/promoter pairs predicted from co-expression across tissues^35^. The LINSIGHT scores for these paired enhancers and promoters are weakly but significantly correlated (Figure 5b), indicating that the same types of evolutionary pressures tend to act at both members of each pair. Together, these results indicate that the evolutionary constraints on enhancers are dependent on several factors, including their degree of tissue specificity, the particular tissues in which they are active, and the evolutionary constraints associated with their target promoters.

**Figure 5:**
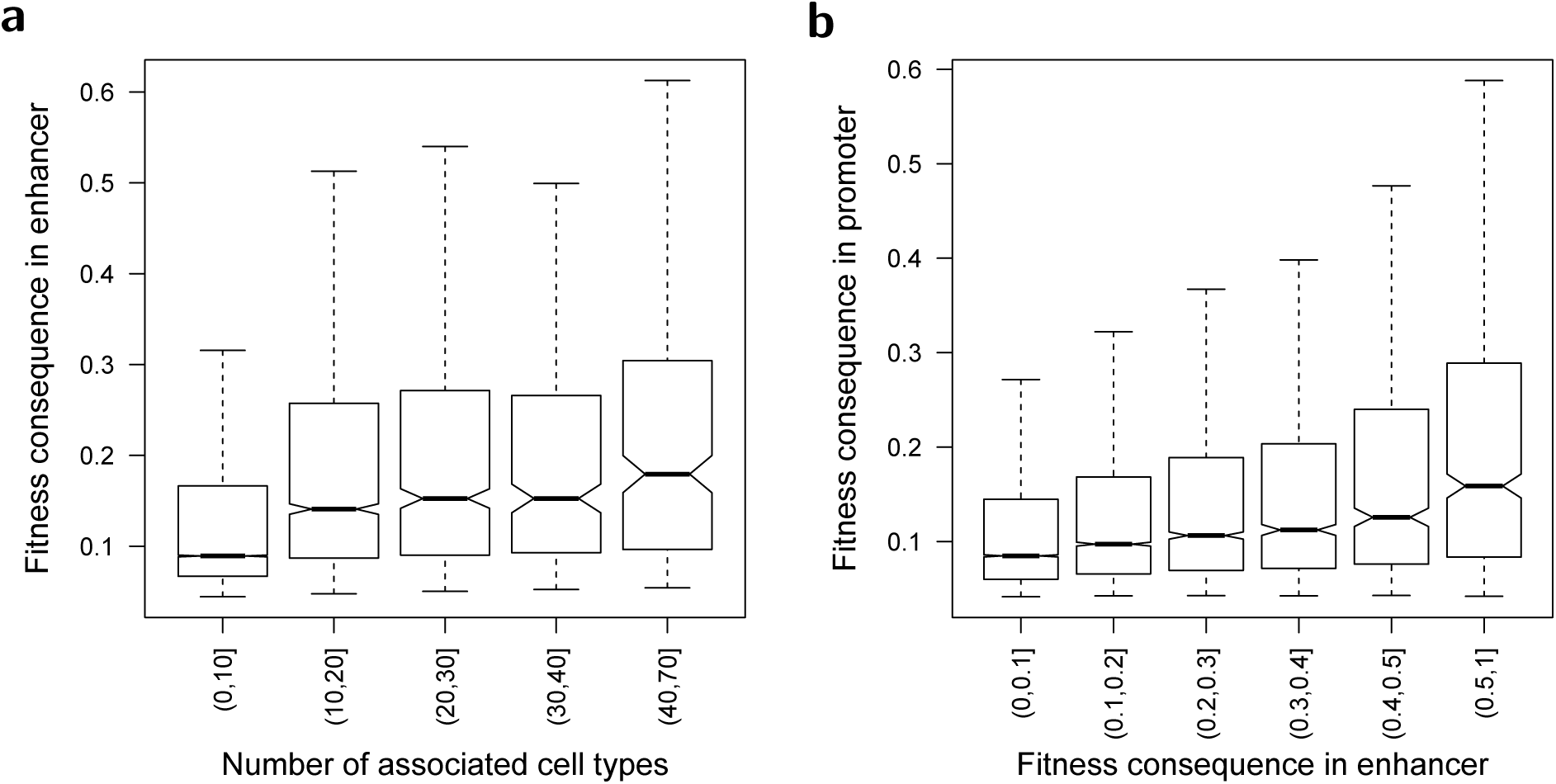
Evolutionary constraints on enhancers. (**a**) Fitness consequences of mutations in enhancers (measured by average LINSIGHT score) is positively correlated with the number of cell types in which each enhancer is active (Spearman's rank correlation coefficient *ρ* = 0.284; *p* < 10^−15^). Results are shown for 29,303 enhancers in 69 cell types. (**b**) Fitness consequences of mutations in enhancers is positively correlated with fitness consequences of mutations in associated promoters (Spearman's rank correlation coefficient *ρ* = 0.150; *p* < 10^−15^). Results are shown for 25,067 enhancer-promoter pairs.

## Discussion

As sequencing costs fall and appreciation for regulatory variation grows, whole genome sequencing is rapidly supplanting exome sequencing as the primary technique for identifying and characterizing genetic variants that have phenotypic consequences. Hence, there is an increasing need for computational methods that can effectively prioritize noncoding variants based on their likelihood of phenotypic importance. In this paper, we address this problem with a new computational method, called LINSIGHT, that combines the evolutionary model of our previously developed INSIGHT method with a generalized linear model for functional genomic data and genome annotations, resulting in substantially improved scalability, resolution, and power. We have generated LINSIGHT scores across the human genome, making use of a large collection of publicly available population, comparative, and functional genomic data, and we find the scores to be consistent with previously available scoring in many respects, but to improve on them in others. For example, they have better power than both pure conservation-based methods and methods based on genomic polymorphism for identifying known transcription factor binding sites. They are competitive with modern machine-learning methods, such as DeepSEA^16^, that are specifically designed for binding site identification. On the task of identifying human disease-associated variants from the HGMD and ClinVar databases, LINSIGHT offered the best performance of several methods we tested, across a range of types of variants and test designs. Importantly, LINSIGHT requires no training set of known regulatory or disease variants and therefore is expected to have better generalization properties and fewer biases than “supervised” machine-learning classifiers or sequence-and motif-based predictors (see Introduction). This improved generalization was evident in the improved performance of LINSIGHT across tests sets compared with GWAVA.

In conceptual terms, the new LINSIGHT method is closely related to our previous fitCons method^3^, with the primary difference being that LINSIGHT pools data across sites implicitly through the use of its linear-sigmoid model, whereas fitCons pools data by explicitly clustering sites according to discretized functional genomic signatures. In effect, LINSIGHT trades the restrictions of a linearity assumption for the benefits of computational speed, a reduced parameterization, and scalability to very large numbers of genomic features. Notably, the new model design also has a number of important side benefits. First, it avoids the need for discretization of the genomic features. In addition, as the number of features grows larger, the genomic resolution of the scores naturally becomes much finer, approaching the nucleotide-level resolution of conservation scores. Finally, the linear-sigmoid model can naturally be extended to a “deep” neural network through the addition of hidden layers. While it remains to be seen how much this extension will help in practice, in principle it can address the types of nonlinearity and interactions between features that have been observed in this setting (for examples, see references [3] and [42]), and it may therefore improve estimates of the fitness consequences of noncoding mutations.

Our approach to characterizing noncoding variants is based on the premise that natural selection in the past, at individual nucleotide sites, provides useful information about phenotypic importance in the present. This assumption clearly will not hold in all cases. For example, variants that increase the risk for post-reproductive diseases or that influence phenotypes dependent on features of the modern environment (such as smoking, industrial chemicals, or abundant high-calorie food) will not necessarily show signs of historical purifying selection. In addition, traits dependent on highly epistatic loci or on the aggregate contributions of large numbers of loci may have difficult-to-detect marginal contributions to fitness at individual nucleotides. Nevertheless, our results indicate that the evolution-based approach is useful for many phenotypes of interest. Furthermore, it is important to bear in mind that experimental approaches for identifying gene regulatory mutations also have limitations. For example, methods that depend on reporter gene assays may not adequately consider the true genomic context and cell-type-dependence of a mutation, and methods that depend on an RNA expression readout may not consider post-transcriptional or post-translational influences. Evolution-based methods have the important advantage of measuring the importance of genetic variants in real organisms in their natural environments over many generations. Thus, we expect that these methods will remain a powerful, complementary tool for characterizing regulatory sequences, even as experimental methods improve.

In our previous work, we avoided considering evolutionary conservation as a genomic feature, instead making a clear distinction between features based on functional genomics and an objective function based on patterns of genetic variation (via INSIGHT)^3^. In this work, we found that we could improve our prediction performance substantially by relaxing this distinction and including conservation scores as inputs to the model (see also references [13, 16, 18]). Thus, despite the limitations of conventional conservation scores—such as their sensitivity to alignment error and evolutionary turnover—they appear to be among the most informative features about the recent natural selection measured by the INSIGHT model. We attribute their value in this setting to the freedom of LINSIGHT to find a weighted combination of conservation scores, functional genomic data, and annotations that is most informative about recent selection. The value of conservation scores may increase further in an extension to a deep neural network, because conservation is likely to be more informative in the presence of some combinations of genomic features than others. For example, evolutionary conservation on the timescale of mammalian evolution is likely to be more informative about recent selection in protein-coding genes, promoters, and splice sites than in enhancers, which exhibit more turnover^43^.

Using LINSIGHT, we examined the influence of negative selection on enhancers, considering the relationships between constraint on enhancers and numbers of active cell types, tissue of activity, and constraint at associated promoters. LINSIGHT is potentially useful for addressing these questions because it should be much more robust to evolutionary turnover than conventional conservation-based methods, and some classes of enhancers are known to turn over more quickly than others^43^. We found that, in general, the trends in constraint at enhancers parallel those previously reported for protein-coding genes. For example, constraint increases with breadth of activity across cell types and decreases in tissues associated with adaptation, such as olfactory regions, the immune system, and male reproduction. Constraint also appears to be correlated at enhancer/promoter pairs. These observations about the specific ways in which evolutionary constraints on enhancers depend on genomic context may be useful in improving the prediction power for the fitness consequences of noncoding mutations.

As has been suggested for protein-coding genes^38^, it seems plausible that the positive correlation between the strength of constraint and the number of active cell types can be explained by pleiotropy: enhancers active in more cell types are more likely to participate in multiple regulatory networks, perhaps with distinct roles involving the binding of different factors and/or the use of different binding sites within each enhancer. As a result, they may be subject to greater constraint. Notably, some of the other explanations offered for a similar correlation between breadth of expression and constraint in protein-coding genes—such as selection for translational efficiency^44,45^ or against misfolding^39^—are not relevant in the case of enhancers. Nevertheless, many open questions remain about the influences of constraint on enhancers, and it will be important to examine these questions further in light of rapidly improving enhancer annotations, data describing enhancer-promoter interactions^46–48^, and observations of complex evolutionary behavior at enhancers^49^.

**URLs.** UCSC Genome Browser, http://genome.ucsc.edu/; Cold Spring Harbor Laboratory Mirror of UCSC Genome Browser, http://genome-mirror.cshl.edu/; Complete Genomics human variation data, http://www.completegenomics.com/public-data/69-Genomes/; SPIDEX database, http://www.deepgenomics.com/spidex/.

## Acknowledgments

We thank other members of the Siepel Laboratory for helpful discussions. This research was supported by US National Institutes of Health (National Institute of General Medical Sciences, NIGMS) grant GM102192. The content is solely the responsibility of the authors and does not necessasrily represent the official views of the US National Institutes of Health.

## Author Contributions

YFH and AS conceived and designed the study. YFH designed and implemented the LINSIGHT method. YFH and BG analyzed the data. AS supervised the research. YFH and AS wrote the manuscript with review and feedback from BG.

## Online Methods

### Genomic features

The genomic features used by LINSIGHT can be divided into three categories: conservation scores, predicted binding sites, and regional annotations (Table 1 and Supplementary Table 2). Conservation scores included phyloP scores^26^, phastCons elements^4^, SiPhy omega elements^50,51^, and CEGA elements^52^. Except for SiPhy, each score type was represented by multiple data tracks—for example, phastCons tracks for vertebrate, mammalian, and primate alignments (Supplementary Table 2). Predicted binding sites included transcription factor binding sites (TFBS) and RNA binding sites. Predicted TFBSs were obtained from the conserved TFBS track in the UCSC Genome Browser^53^, the rVISTA database^54^, SwissRegulon^55^, FunSeq2^20^, and the Ensembl Regulatory Build^27^. RNA binding sites include splice sites predicted by SPIDEX^56^ and miRNA target sites predicted by TarBase^57^. The regional annotations were based a variety of sources, including ChIP-seq and RNA-seq data from the ENCODE^11^ and Roadmap Epigenomics^12^ projects, enhancers from FANTOM5^58^, predicted distal regulatory modules from FunSeq2^20^, and the distances to nearest TSSs based on GENCODE gene models^59^. All features and the resulting LINSIGHT scores were expressed in genomic coordinates for the hg19 assembly of the human genome.

### Polymorphism and divergence data

The polymorphism and divergence data used by the INSIGHT component of the LINSIGHT model were borrowed from previous analyses^3,21,22^. Briefly, we obtained human single nucleotide polymorphisms from high-coverage genome sequences for 54 unrelated individuals from the “69 Genomes” dataset from Complete Genomics, eliminating nucleotide sites with more than two alleles. Outgroup alleles were defined by the aligned chimpanzee, orangutan, and rhesus macaque reference genomes from UCSC. Several filters were applied to these data to reduce technical errors from alignment, sequencing, and genotype inference; for example, we removed simple repeats, recent transposable elements, recent segmental duplications, and regions not in syntenic alignment across primates^22^. Putatively neutral regions were identified by starting with all aligned regions, then removing coding sequences, conserved non-coding sequences, and their proximal flanking regions. These regions were used to estimate neutral divergence and polymorphism rates in the human lineage in a block-wise manner across the genome, to account for local variation in mutation rates^21^. To allow for uncertainty in the human-chimpanzee most recent common ancestor (MRCA), we integrated over a distribution of ancestral alleles inferred after fitting a standard phylogenetic model to the outgroup sequences^21^.

### Fitting the LINSIGHT model to the data

The weights for all genomic features were estimated by approximately maximizing the log likelihood of the INSIGHT model with respect to our genome-wide data set. We began by considering all genomic positions not excluded by our data-quality filters. Because our focus was on noncoding regions, we additionally excluded coding regions annotated by GENCODE (release 19). Instead of a traditional “batch” learning algorithm, which would require either storing all data in memory or reading it from disk many times, we used an “online” stochastic gradient descent algorithm^60^. The algorithm processed the genome in “minibatches” of 100 nucleotides, each time updating the parameter vector in the direction of the gradient of the log likelihood function, with learning rates of 0.001 and 0.01 for *ρ* and *γ*, respectively. Gradients were computed analytically, by propagating partial derivatives through the linear-sigmoid component of the model using the chain rule (back-propagation). The learning procedure was truncated after 20 passes through the data set. The entire process took less than one day on a desktop computer. The genome-wide LINSIGHT scores are available from the Cold Spring Harbor Laboratory mirror of the UCSC Genome Browser (hg19 assembly).

### Comparison with other methods

Our benchmarking scheme for prioritization of disease-associated variants closely followed the one introduced in ref. [13]. The HGMD and ClinVar noncoding disease variants and three sets of negative controls were obtained from this study. The negative controls consisted of: (1) a randomly selected subset of human common variants (unmatched); a subset of human common variants matched to the disease variants by distance-to-nearest-TSS (matched TSS); and (3) a subset of human common variants required to be in the same local genomic region as the matched disease variants (matched region). The two matched sets account for the enrichment of known disease variants near coding genes. For comparison, we downloaded precomputed CADD^18^ (v1.3), GWAVA^13^ (v1.0), FunSeq2^20^ (v2.1.0), and Eigen^32^ (Oct. 11, 2015) scores from the source websites. GWAVA scores based on training with variants matched by distance-to-nearest-TSS were used in all comparisons^13^. In addition, we obtained mammalian phyloP^26^ scores based on the 46-way whole-genome alignment for hg19 from the UCSC Genome Browser^53^. The DeepSEA scores of the disease variants and their negative controls were computed using the online DeepSEA web service^16^ on Jan 14, 2015. Note that two of the methods considered, CADD and DeepSEA, provide allele-specific predictions, whereas the other methods assign identical scores to all alternative variants. When evaluating CADD and DeepSea on the ClinVar data set, we used the score corresponding to the annotated disease-associated allele. When evaluating these methods on the HGMD data set, however, no disease-associated allele was provided, so we used the maximum score for the three alternative alleles.

### Classification of disease-associated variants by genomic location

For analyses that considered the genomic locations of disease-associated variants, we divided the variants in the HGMD and ClinVar databases into four categories based on their locations relative to gene models from GENCODE (release 19). These categories were: (1) “promoter” variants, located within 1 kb upstream of the 5′-most annotated transcription start site of any protein-coding gene; (2) “splicing” variants, located within 20 bp of any annotated splice junction; (3) “UTR” variants, located within the annotated 5′ or 3′ UTR of any protein-coding gene; and (4) all “other” variants. Each variant was assigned to the first category whose criteria it fulfilled in the order *splicing > UTR > promoter > other*.

### Quantification of the contributions of genomic feature classes

We measured the relative contributions of the conservation scores, predicted binding sites, and regional annotations by removing all features of each class (see Table 1), retraining the LINSIGHT model without those features, and evaluating the AUC of the reduced model. The *contribution* of each class of features was defined as the AUC for the full model minus the AUC for the reduced model. Notice that, while this difference in AUCs is generally positive, it may be negative due to stochastic effects. This analysis was performed on a merged set of HGMD and ClinVar variants, separately for each of the four genomic location labels defined above.

### Analysis of evolutionary constraints on enhancers

To study evolutionary constraints on enhancers, we used the comprehensive atlas of human enhancers across cell types based on enhancer RNAs (eRNAs) that was recently provided by the FANTOM5 project^58^. The evolutionary constraint for each enhancer was quantified by taking the average LINSIGHT score across all nucleotide sites in the enhancer. We examined the relationship between this measure of constraint and the number of cell types in which each enhancer was active (according to a detectable eRNA signature). We also defined a subset of enhancers as tissue-specific, based on apparent activity in only a single tissue type, and examined the relationship between tissue of activity and degree of constraint. Finally, we obtained putative enhancer-TSS pairs (based on correlated patterns of expression across tissues) from the FANTOM5 website, and examined the correlation in constraint at the enhancer and promoter in each pair, defining the promoter as the 1 kb region upstream of the TSS. In cases where an enhancer was associated with multiple TSSs, the TSS with highest correlation coefficient was used as the matched TSS.

## Supplementary material

### Details of the LINSIGHT model

#### Three modes of evolution

LINSIGHT infers the strength of natural selection based on an evolutionary model closely related to the INSIGHT model^1,2^. The key idea of this model is that natural selection and neutral evolution generate distinct patterns of variation in divergence and polymorphism data. In LINSIGHT, we assume that the evolutionary process at a noncoding site occurs by one of three modes: neutral drift (neut), weak negative selection (WN), and strong negative selection (SN). Furthermore, the evolutionary mode of a site does not change over time along the human lineage (since the human/chimpanzee divergence). These assumptions are the same as those of INSIGHT, except in this case we exclude the possibility of positive selection, which turns out to have negligible importance when estimating genome-wide probabilities of fitness consequences^3^.

LINSIGHT is able to distinguish among these three modes of evolution because they have different effects on divergence and polymorphism patterns. In particular, as in INSIGHT, we assume that mutations in strongly negatively selected sites are immediately removed from the population and cannot segregate or fix in the human population. In addition, we assume that mutations in weakly negatively selected sites can segregate at low frequencies but cannot reach high frequencies or fix in the human population. In contrast, mutations in neutral sites can segregate at both low and high frequencies and can fix in the human population.

#### Polymorphism and divergence data

LINSIGHT gains power by combining the signatures of natural selection from both polymorphisms within the target species and divergences between species. As in INSIGHT, each site in the human genome is first classified as monomorphic (*M*), polymorphic with a low-frequency minor allele (*L*), or polymorphic with a high-frequency minor allele (*H*). The minor allele frequency (MAF) at site *i* is denoted by *m_i_*, and its low-dimensional, categorical summary (*M/L/H*) is denoted by *Y_i_*. Sites without alternative alleles, i.e., with *m_i_* = 0, are defined as *M* sites. *L* and *H* sites are distinguished by a MAF threshold *f*, i.e., *Y_i_* = *L* if 0 < *m_i_* < *f* and *Y_i_* = *H* if *f* ≤ *m_i_* ≤ 0.5. It has been shown that this low dimensional summary of the allele frequency data helps to make the model robust to complex demographic histories, at little cost in inference power^1^. As in our previous work, we set *f* to be equal to 0.15. The polymorphism data at site *i* in the human population is summarized as 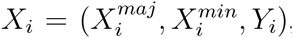, in which 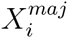 is the observed major allele, 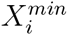 is the observed minor allele, and *Y_i_* ∈ {*M, L, H*} is the minor allele frequency class. In the whole genome alignment of human and other primates, the aligned bases in the primate outgroup at site *i* are denoted by the vector **O***_i_*. See references [1] and [2] for further details.

#### Parameters in the LINSIGHT model

As in INSIGHT, inferences of natural selection depend on the contrast between a model that allows for selection and a neutral model. LINSIGHT uses the same neutral model as INSIGHT ^1^. Briefly, for each site *i* in the human genome, two neutral parameters are specified: *λ_i_* and *θ_i_*. The parameter *λ_i_* denotes the block-wise neutral substitution rate associated with site *i* in the human lineage after the human-chimpanzee divergence. Similarly, *θ_i_* denotes the block-wise neutral polymorphism rate associated with site *i*. In addition, there is a set of three global parameters, denoted *β* = (*β*_1_, *β*_2_, *β*_3_), that describe the genome-wide proportions of low, intermediate, and high frequency derived neutral alleles, respectively. Note that *β* is defined based on the frequencies of derived alleles, rather than the MAF, and therefore depends on the ancestral allele. The estimation procedure for *β* integrates over possible values of the ancestral allele. Together, we denote the neutral parameters by *ζ_i_* = (*λ_i_, θ_i_, β*). These parameters were reused from our previous work. See references [1] and [2] for further details.

Similar to INSIGHT, LINSIGHT summarizes the influence of negative selection at site *i* using two parameters: *ρ_i_* and *γ_i_*. (A third selection parameter considered by INSIGHT, *η_i_*, is not needed because of the omission of positive selection.) The parameter *ρ_i_* describes the probability that site *i* is under selection (WN or SN). The parameter *γ_i_* represents the relative rate of low-frequency derived alleles at sites under selection. These two parameters together describe whether and how the evolutionary process at site *i* diverges from that at the flanking neutral sites.

The parameter *ρ_i_* is the main focus of our analysis. This parameter can be interpreted as the probability that a mutation at that site will have fitness consequences. Nucleotide-specific estimates of this parameter are therefore referred to as fitness-consequence scores, or simply, as LINSIGHT scores. The parameter *γ_i_* is influenced by both weak and strong selection and is more difficult to interpret. This parameter is used for fitting the model but is ignored in our subsequent analyses. Note that the definition of *γ_i_* here is slightly different from its definition in the INSIGHT model (denoted here by 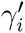), which was defined as the relative rate of polymorphisms. There is a simple linear relationship between *γ_i_* and 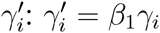. This redefinition of *γ_i_* was needed to ensure that *ii* its range would match that of the sigmoid function (0, 1).

Because the information about natural selection at each individual nucleotide is very limited, some means for pooling statistical information across sites is needed. In LINSIGHT, this pooling is achieved by a generalized linear regression model. We assume that the selection parameters at each focal site can be predicted by a linear function of local genomic features, together with a nonlinear “link” function. Specifically, if **D***_i_* is a column vector of genomic features associated with site *i*, LINSIGHT assumes that,

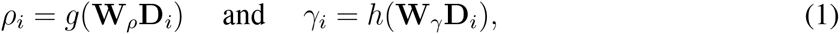

where the row vectors **W***_ρ_* and **W***_γ_* represent the weights of genomic features with respect to *ρ* and *γ*, respectively, and *g*() and *h*() are nonlinear link functions. Following the common practice in machine learning, we assume that the first element of **D***_i_* is always equal to 1 and the corresponding weights in **W***_ρ_* and **W***_γ_* represent the intercepts for the respective linear functions. For the link functions *g*() and *h*(), we use two sigmoid functions,

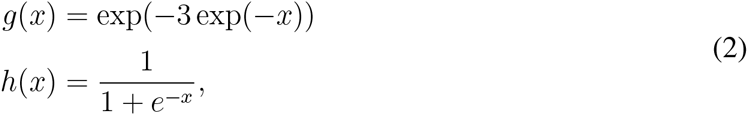

where *g*(*x*) is known as the Gompertz sigmoid function^4^ and is used here to avoid saturation at small values of *x*. Thus, the linear combinations of genomic features are mapped to the range between 0 and 1, separately for each of the two parameters at each nucleotide site. Because **W***_ρ_* and **W***_γ_* are shared by all sites, this strategy allows sharing of statistical strength across the genome.

#### Likelihood function

To estimate the **W***_ρ_* and **W***_γ_* vectors (the free parameters of the LINSIGHT model), and hence to obtain the nucleotide-specific LINSIGHT scores, an objective function for optimization is needed. For this purpose, we use the likelihood function defined for INSIGHT, with minor changes to accommodate the omission of positive selection (*η* parameter). Maximization of this likelihood function will therefore allow for maximum likelihood estimation of **W***_ρ_* and **W***_γ_*. The combination of the adapted INSIGHT model and our generalized linear model is depicted in Figure 1 of the main text. In the following discussion, we briefly detail key features of this model, focusing on differences from the original INSIGHT model.

As in INSIGHT, we introduce two latent variables, *Z_i_* and *A_i_*, representing the alleles at site *i* in the most recent common ancestors of human-chimpanzee and the human individuals, respectively. The evolutionary trajectory of site *i* in the human lineage can then be denoted by (*Z_i_, A_i_, X_i_*). Another binary latent variable, *S_i_*, is introduced to represent whether or not site *i* is under negative selection. The likelihood function for site *i* thus requires summing over all possible configurations of *S_i_*, *Z_i_*, and *A_i_*. Assuming independence across sites, the likelihood function for the whole genome is given by:

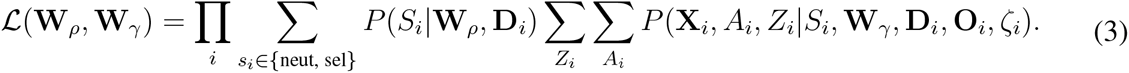

Here, *P* (*S_i_*|**W***_ρ_*, **D***_i_*) represents the *prior* probability that site *i* is under selection conditional on genomic features **D***_i_* and weights **W***_ρ_*. Based on equation 1, this prior probability is given by

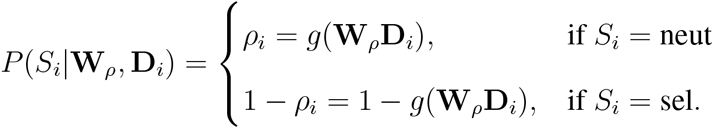

The other probability in equation 3, *P* (**X***_i_, A_i_, Z_i_|S_i_*, **W***_γ_*, **D***_i_*, **O***_i_, ζ_i_*) representing the probability of the evolutionary trajectory (*Z_i_, A_i_, X_i_*), can be factorized as follows based on the conditional independence assumptions of the model (as in INSIGHT):

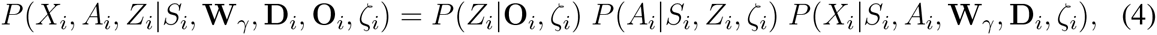

where *P* (*Z_i_|***O***_i_, ζ_i_*) represents the prior distribution of the deep ancestral allele *Z_i_*, *P* (*A_i_*|*S_i_, Z_i_, ζ_i_*) represents the generation of fixed mutations (substitutions) along the human lineage, and *P* (*X_i_*|*S_i_, A_i_*, **W***_γ_*, **D***_i_, ζ_i_*) represents the generation of polymorphisms in the human population. As in INSIGHT, *P* (*Z_i_|***O***_i_, ζ_i_*) is estimated using a standard phylogenetic model and *P* (*A_i_|S_i_, Z_i_, ζ_i_*) is modeled by an approximate Jukes-Cantor substitution model^5,6^. Here,

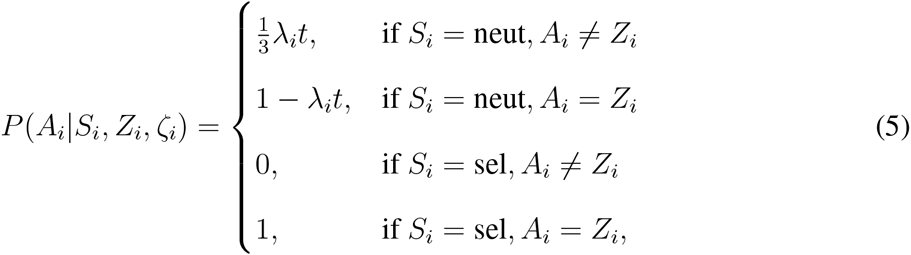

where *t* represents the evolutionary time between the human-chimpanzee MRCA and the human population MRCA. Note that, unlike in INSIGHT, when site *i* is under selection, no substitution is possible because positive selection is prohibited in LINSIGHT. As in INSIGHT, however, recurrent substitutions at the same site are ignored due to the very short divergence time between human and chimpanzee.

The final term in equation 4, *P* (*X_i_|S_i_, A_i_*, **W***_γ_*, **D***_i_, ζ_i_*), represents the generation of (unfixed) polymorphisms in the human population. As in INSIGHT, we assume a simple infinite-sites model for the generation of polymorphisms, in which polymorphisms are generated at rate *θ_i_* (and all three alternative alleles are equally likely), and the allele frequency category (*L* or *H*) is determined by the *β* and *γ_i_* parameters. Specifically, we assume:

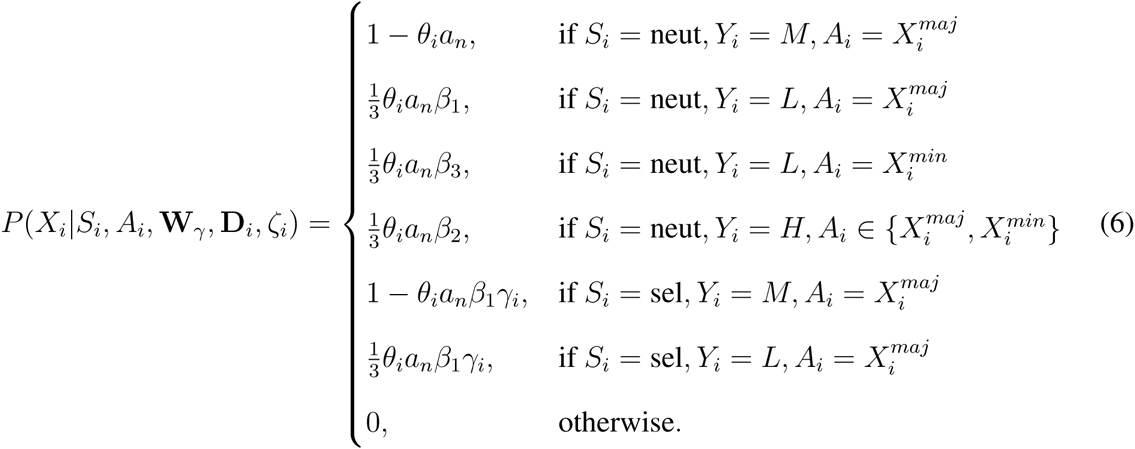

where *n* is the number of haploid genomes sampled at site *i*, 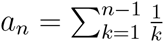 is Watterson's constant^7^, and *γ_i_* is given by equation 1. As in INSIGHT, Watterson's constant *a_n_* can be used to accommodate small amounts of missing data^1^. Notice in equation 6 that polymorphisms in sites under selection (*S_i_* = sel) are only permitted to segregate at low frequencies, consistent with the assumptions of the model. Combining equations 5 and 6 and then summing over possible values of *A_i_*, we obtain the conditional likelihood *P* (*X_i_|S_i_, Z_i_*, **W***_γ_*, **D***_i_, ζ_i_*), which is summarized in Table 1. Finally, the full likelihood (equation 3) is calculated by summing over possible values of *Z_i_*.

---

**Supplementary Table 1:** Conditional distribution table for *P* (*X_i_|S_i_, Z_i_*, **W***_γ_*, **D***_i_, ζ_i_*)

| $S_i$ | $Y_i$ | $Z_i, X_i^{maj}, X_i^{min}$ | $P(X_i\vert S_i, Z_i, \mathbf{W}_\gamma, \mathbf{D}_i, \zeta_i)^{ab}$ |
| --- | --- | --- | --- |
| neut | $M$ | $Z_i = X_i^{maj}$ | $(1 - \lambda_i t)(1 - \theta_i a_n)$ |
| neut | $M$ | $Z_i \neq X_i^{maj}$ | $\frac{1}{3} \lambda_i t (1 - \theta_i a_n)$ |
| neut | $L$ | $Z_i = X_i^{maj}$ | $((1 - \lambda_i t) \beta_1 + \frac{1}{3} \lambda_i t \beta_3) \frac{1}{3} \theta_i a_n$ |
| neut | $L$ | $Z_i = X_i^{min}$ | $((1 - \lambda_i t) \beta_3 + \frac{1}{3} \lambda_i t \beta_1) \frac{1}{3} \theta_i a_n$ |
| neut | $L$ | $Z_i \notin \{X_i^{maj}, X_i^{min}\}$ | $\frac{1}{3} \lambda_i t (\beta_1 + \beta_3) \frac{1}{3} \theta_i a_n$ |
| neut | $H$ | $Z_i \in \{X_i^{maj}, X_i^{min}\}$ | $(1 - \lambda_i t + \frac{1}{3} \lambda_i t) \beta_2 \frac{1}{3} \theta_i a_n$ |
| neut | $H$ | $Z_i \notin \{X_i^{maj}, X_i^{min}\}$ | $\frac{2}{3} \lambda_i t \beta_2 \frac{1}{3} \theta_i a_n$ |
| sel | $M$ | $Z_i = X_i^{maj}$ | $1 - \beta_1 \gamma_i \theta_i a_n$ |
| sel | $M$ | $Z_i \neq X_i^{maj}$ | 0 |
| sel | $L$ | $Z_i = X_i^{maj}$ | $\frac{1}{3} \beta_1 \gamma_i \theta_i a_n$ |
| sel | $L$ | $Z_i \neq X_i^{maj}$ | 0 |
| sel | $H$ | — | 0 |

<sup>a</sup> $\zeta_i = (\lambda_i, \theta_i, \beta)$ represents the set of neutral parameters associated with site $i$ .
<sup>b</sup> $\gamma_i = h(\mathbf{W}_\gamma \mathbf{D}_i)$ represents the relative rate of low-frequency derived alleles.

**Supplementary Table 2:** Genomic features used for LINSIGHT scores for human genome (hg19)

| Feature | Type | Definition | Weight<sup>a</sup> |
| --- | --- | --- | --- |
| phyloP mammal score (46 way) | Numeric | Truncated at 2 to reduce false positive rate | 0.673 |
| phyloP mammal class (46 way) | Binary | Score $\geq 2$ is coded as 1, otherwise 0 | 1.418 |
| phyloP vertebrate score (46 way) | Numeric | Truncated at 2 to reduce false positive rate | 0.709 |
| phyloP vertebrate class (46 way) | Binary | Score $\geq 2$ is coded as 1, otherwise 0 | 1.456 |
| Primate phastCons element (46 way) | Binary | Presence/absence of conserved element | 0.413 |
| Mammal phastCons element (46 way) | Binary | Presence/absence of conserved element | 0.120 |
| Vertebrate phastCons element (46 way) | Binary | Presence/absence of conserved element | 0.227 |
| SiPhy omega element (29 way) | Binary | Presence/absence of conserved element | 0.049 |
| CEGA conserved element (Amniota) | Binary | Presence/absence of conserved element | 0.000 |
| CEGA conserved element (Boreoeutheria) | Binary | Presence/absence of conserved element | 0.149 |
| CEGA conserved element (Euarchontoglires) | Binary | Presence/absence of conserved element | 0.226 |
| CEGA conserved element (Eutheria) | Binary | Presence/absence of conserved element | 0.126 |
| CEGA conserved element (Vertebrata) | Binary | Presence/absence of conserved element | -0.058 |
| UCSC conserved TFBS | Binary | Presence/absence of TFBS | 0.306 |
| TFBS in Ensembl regulatory build | Binary | Presence/absence of TFBS | 0.098 |
| rVISTA TFBS | Binary | Presence/absence of TFBS | 0.232 |
| SwissRegulon TFBS | Binary | Presence/absence of TFBS | 0.179 |
| FunSeq2 (ENCODE) TFBS | Binary | Presence/absence of TFBS | 0.094 |
| TarBase miRNA target | Binary | Presence/absence of miRNA binding | 0.050 |
| Maximum SPIDEX score | Numeric | Truncated at 2 to reduce false positive rate | 0.228 |
| Maximum SPIDEX score class | Binary | Maximum score $\geq 2$ is coded as 1, otherwise 0 | 0.190 |
| Distance to nearest TSS | Numeric | $\frac{10Kb-dist}{10Kb}$ if $dist \leq 10Kb$, otherwise 0 | 0.051 |
| ENCODE (UCSC) FAIRE peak | Binary | Presence/absence of peak | 0.043 |
| ENCODE (UCSC) FAIRE prevalence | Numeric | Number of tissues associated with the peak | -0.076 |
| ENCODE (UCSC) DNase peak | Binary | Presence/absence of peak | 0.015 |
| ENCODE (UCSC) DNase prevalence | Numeric | Number of tissues associated with the peak | -0.004 |
| Predicted distal regulatory module in FunSeq2 | Binary | Presence/absence of cis-regulatory module | 0.062 |
| ENCODE TF ChIP-seq peak (UCSC) | Binary | Presence/absence of peak | 0.006 |
| ENCODE TF ChIP-seq prevalence (UCSC) | Binary | Number of TFs associated with the peak | -0.003 |
| Maximum Roadmap Epigenomics RNA-seq class | Binary | Site with signal is coded as 1, otherwise 0 | -0.002 |
| Maximum Roadmap Epigenomics RNA-seq signal | Numeric | Log transformed signal if exists, otherwise 0 | 0.129 |
| FANTOM5 enhancer | Binary | Presence/absence of FANTOM5 enhancer | 0.028 |
| ENCODE/Roadmap distal DNase peak | Binary | Presence/absence of peak | -0.006 |
| ENCODE/Roadmap distal DNase prevalence | Numeric | Number of tissues associated with the peak | 0.037 |
| ENCODE/Roadmap distal H3K27ac peak | Binary | Presence/absence of peak | 0.007 |
| ENCODE/Roadmap distal H3K27ac prevalence | Numeric | Number of tissues associated with the peak | 0.031 |
| ENCODE/Roadmap distal H3K4me1 peak | Binary | Presence/absence of peak | 0.010 |
| ENCODE/Roadmap distal H3K4me1 prevalence | Numeric | Number of tissues associated with the peak | 0.031 |
| ENCODE/Roadmap distal H3K9 peak | Binary | Presence/absence of peak | 0.024 |
| ENCODE/Roadmap distal H3K9 prevalence | Numeric | Number of tissues associated with the peak | 0.025 |
| ENCODE/Roadmap distal TF ChIP-seq peak | Binary | Presence/absence of binding site | -0.006 |
| ENCODE/Roadmap proximal DNase peak | Binary | Presence/absence of peak | 0.043 |
| ENCODE/Roadmap proximal DNase prevalence | Numeric | Number of tissues associated with the peak | -0.061 |
| ENCODE/Roadmap proximal H3K4me3 peak | Binary | Presence/absence of peak | 0.024 |
| ENCODE/Roadmap proximal H3K4me3 prevalence | Numeric | Number of tissues associated with the peak | -0.040 |
| ENCODE/Roadmap proximal H3K9ac peak | Binary | Presence/absence of peak | 0.015 |
| ENCODE/Roadmap proximal H3K9ac prevalence | Numeric | Number of tissues associated with the peak | -0.039 |
| ENCODE/Roadmap proximal TFBS | Binary | Presence/absence of binding site | 0.009 |

<sup>a</sup>Estimated weight for $\rho$ after genome-wide training. Weights are directly comparable because all covariates are rescaled to a range between 0 and 1 before training.

**Supplementary Figure 1:**
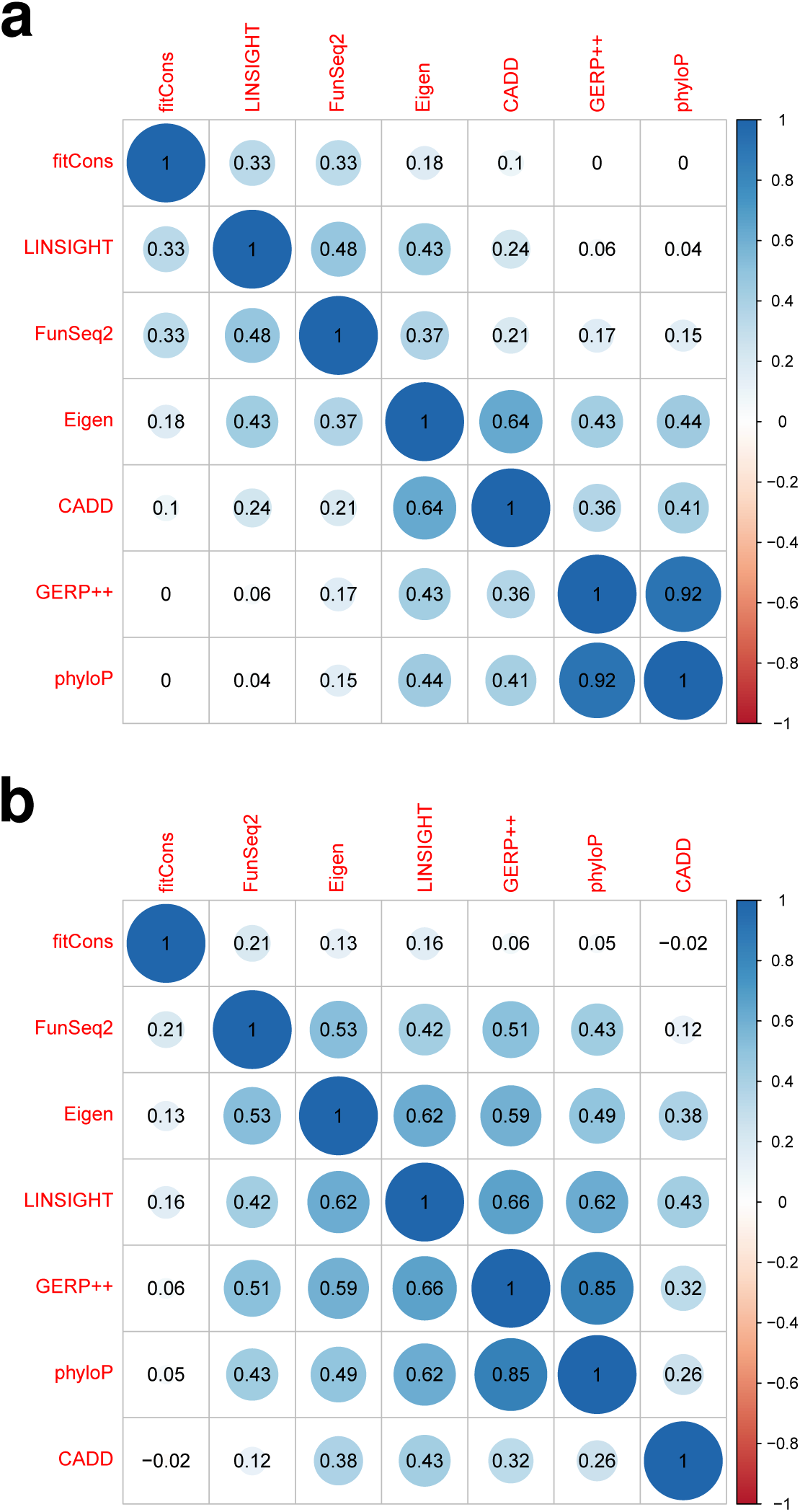
Spearman's correlation coefficients (*ρ*) for all pairs of scores considered. **(a)** Correlation at all scored genomic positions. **(b)** Correlation at sites in mammalian phastCons elements.

**Supplementary Figure 2:**
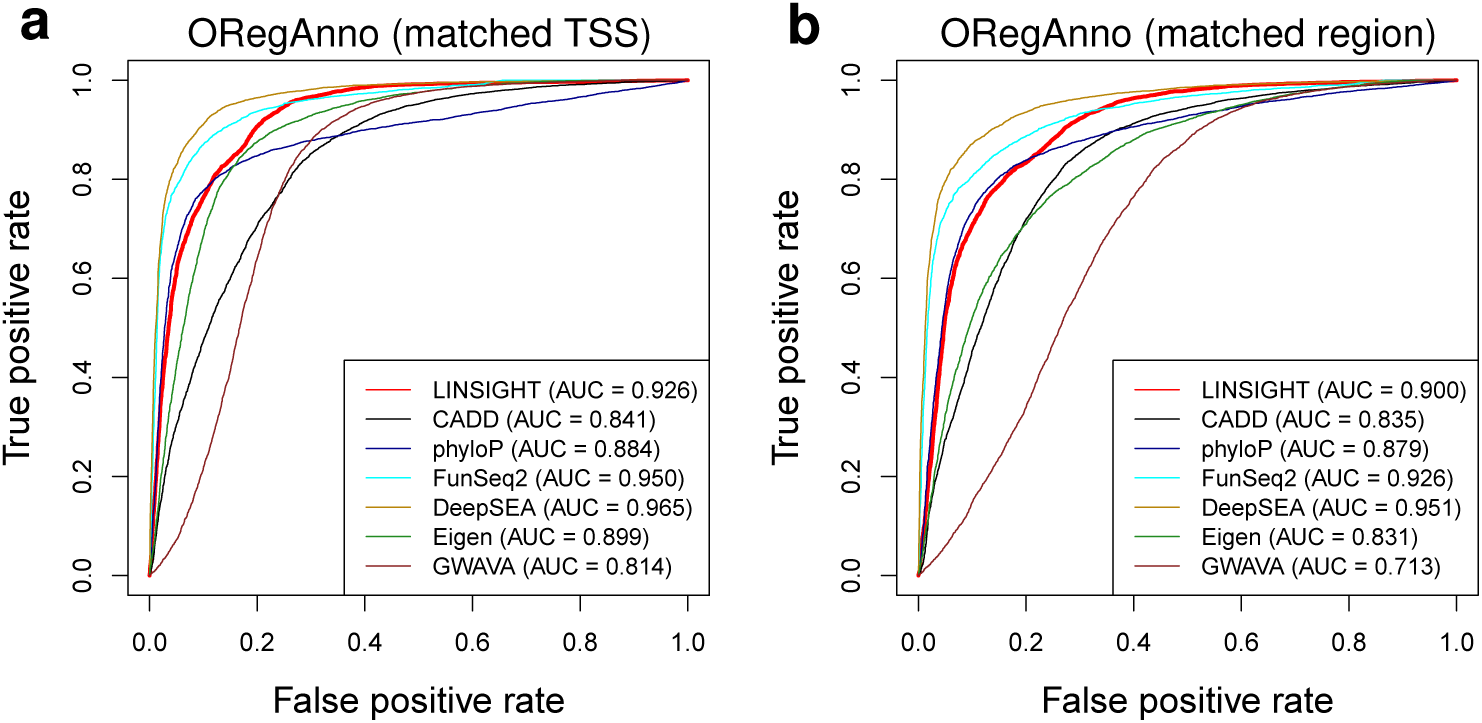
Prediction power of various computational methods for distinguishing predicted transcription factor binding sites (TFBS) from likely non-TFBSs, described as receiver operating characteristic (ROC) curves. Results are shown for the **(a)** “matched TSS” and **(b)** “matched region” schemes for pairing positive and negative examples (see Methods). We considered all TFBSs in the ORegAnno database^8^ that were associated with the hg19 assembly, merging overlapping binding sites (7,369 TFBSs in total).

**Supplementary Figure 3:**
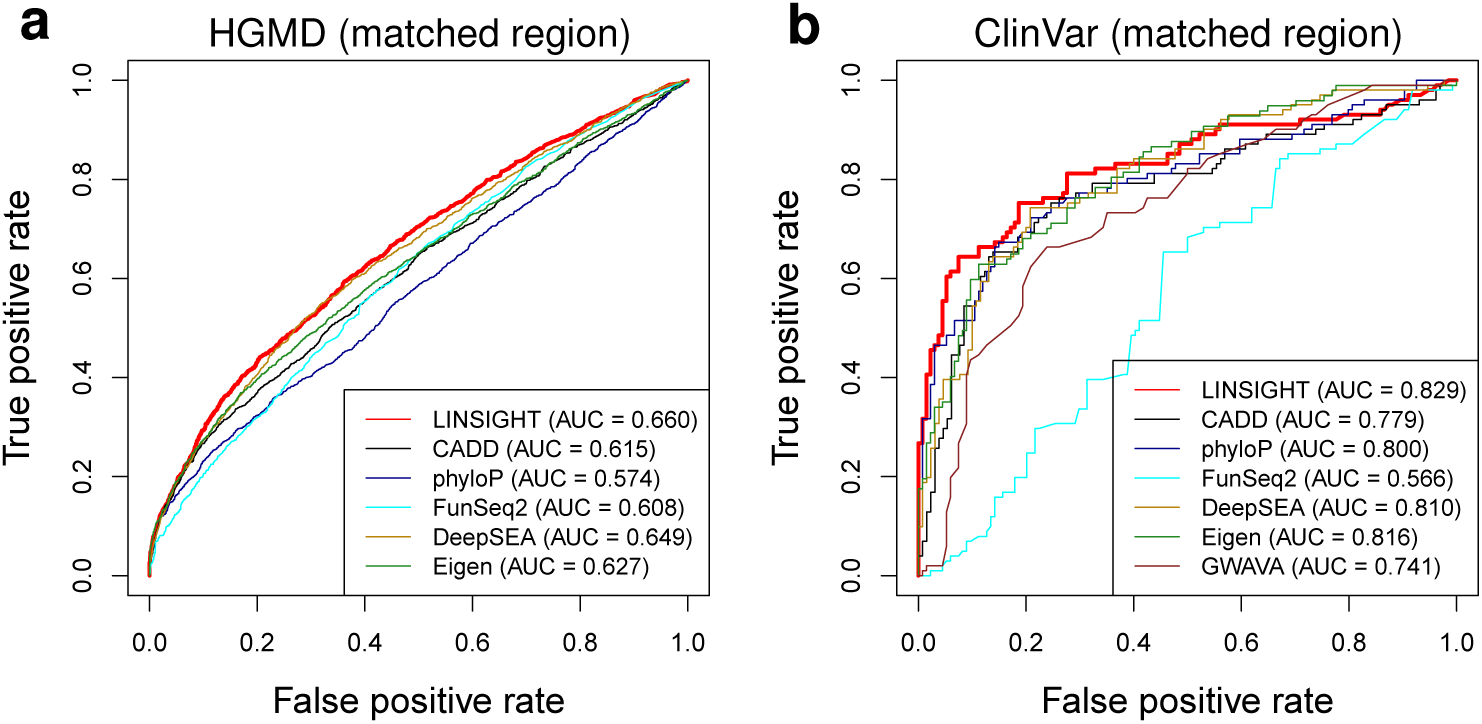
Prediction power of various computational methods for distinguishing disease-associated noncoding variants from variants not likely to have phenotypic effects. Plots are the same as those in Figure 3 except for the use of the region-based matching scheme for positive and negative examples (see Methods). Plots describe positive examples from **(a)** HGMD and **(b)** ClinVar.

**Supplementary Figure 4:**
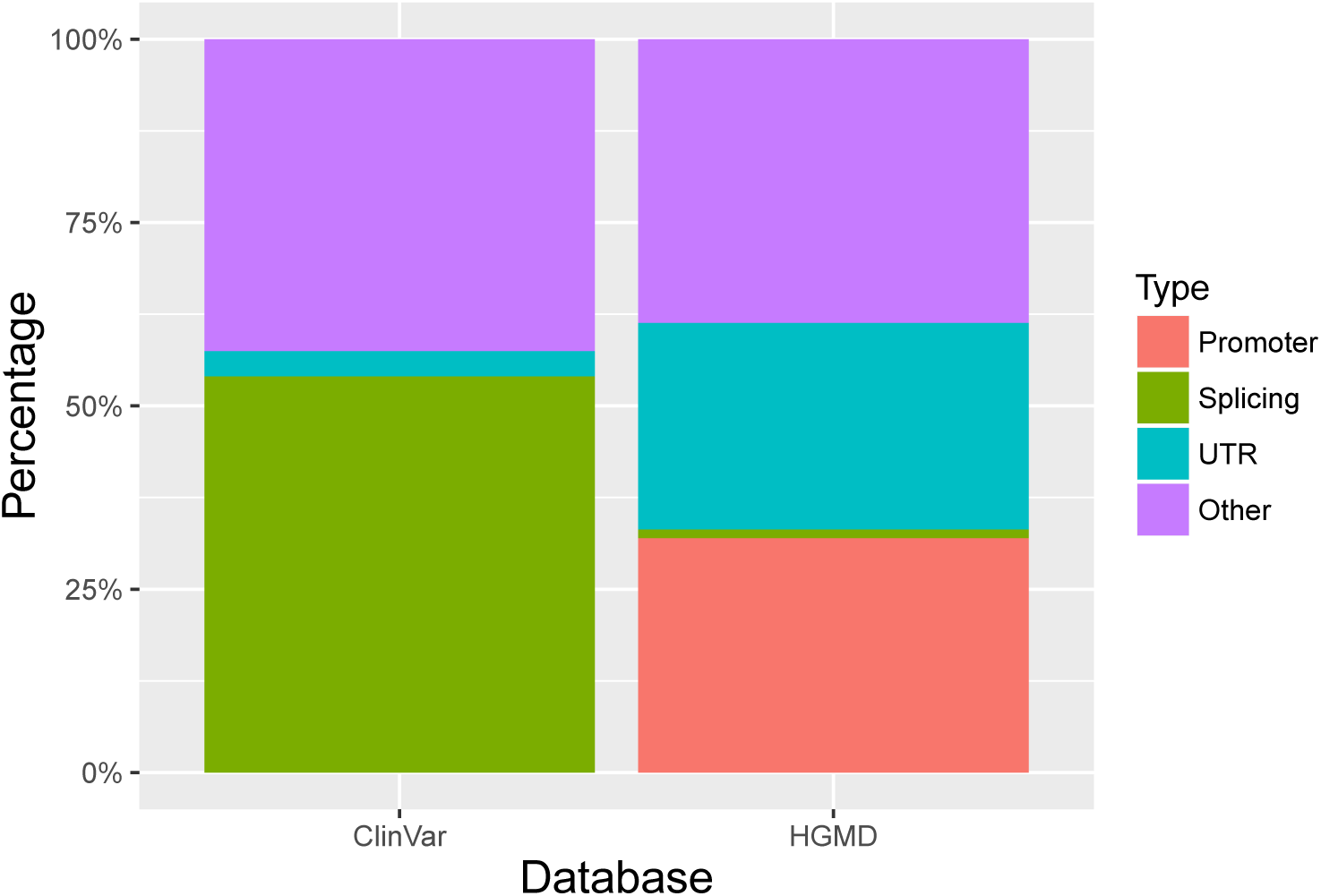
Genomic distributions of noncoding disease variants in the ClinVar (left) and HGMD (right) data sets. See Methods for definitions of Promoter, Splicing, UTR, and Other genomic regions.

**Supplementary Figure 5:**
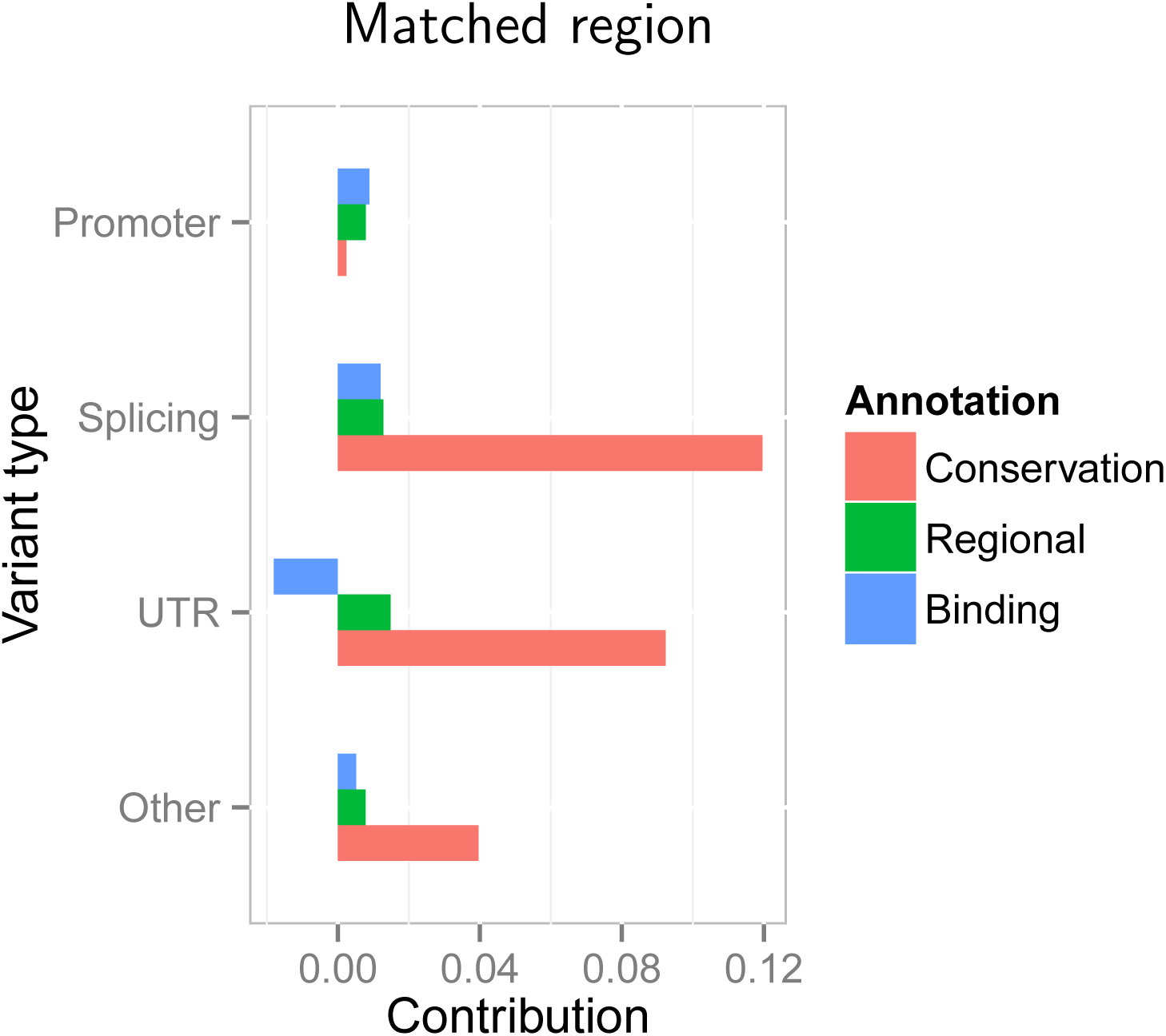
Contributions of various genomic features to the identification of disease-associated variants from HGMD and ClinVar. Plot is the same as those in Figure 4 except for the use of the region-based matching scheme for positive and negative examples (see Methods).

**Supplementary Figure 6:**
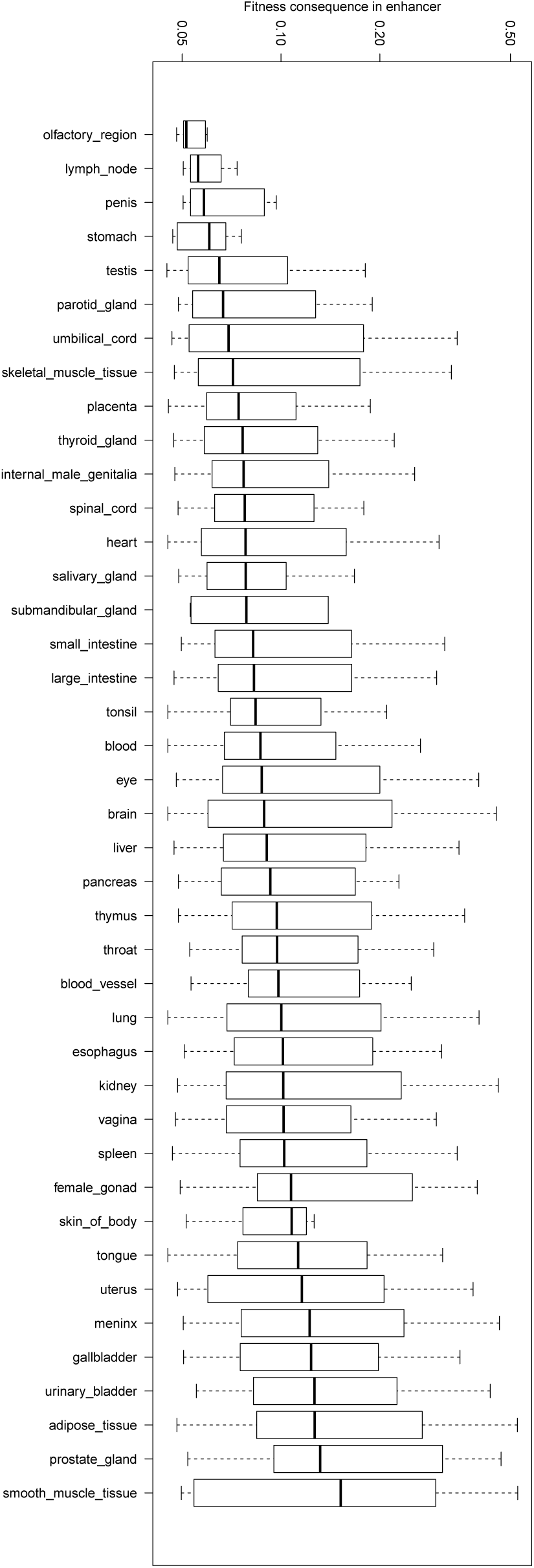
Distribution of fitness consequences of mutations at 8,082 tissue-specific enhancers (measured by average LINSIGHT score per enhancer) across 41 tissue types.

